# Human sperm TMEM95 binds eggs and facilitates membrane fusion

**DOI:** 10.1101/2022.06.10.495573

**Authors:** Shaogeng Tang, Yonggang Lu, Will M. Skinner, Mrinmoy Sanyal, Polina V. Lishko, Masahito Ikawa, Peter S. Kim

## Abstract

*Tmem95* encodes a sperm acrosomal membrane protein, whose knockout has a male-specific sterility phenotype in mice. How TMEM95 plays a role in membrane fusion of sperm and eggs has remained elusive. Here, we utilize a sperm penetration assay as a model system to investigate the function of human TMEM95. We show that human TMEM95 binds to hamster egg membranes, providing evidence for a TMEM95 receptor on eggs. Using X-ray crystallography, we reveal an evolutionarily conserved, positively charged region of TMEM95 as a putative receptor-binding surface. Amino-acid substitutions within this region of TMEM95 ablate egg-binding activity. We identify monoclonal antibodies against TMEM95 that reduce the number of human sperm fused with hamster eggs in sperm penetration assays. Strikingly, these antibodies do not block binding of sperm to eggs. Taken together, these results provide strong evidence for a specific, receptor-mediated interaction of sperm TMEM95 with eggs and suggest that this interaction may have a role in facilitating membrane fusion.

**Significance statement:** Membrane fusion of sperm and eggs is pivotal in sexual reproduction. *Tmem95* knockout mice show male-specific sterility, but it was unknown how sperm TMEM95 facilitates membrane fusion with eggs. We show here that human TMEM95 binds eggs. Our crystal structure of TMEM95 suggests a region where this binding may occur. We develop monoclonal antibodies against TMEM95 that impair sperm-egg fusion but do not block sperm-egg binding. Thus, we propose that there is a receptor-mediated interaction of sperm TMEM95 with eggs, and that this interaction may have a direct role in membrane fusion. Our work suggests avenues for the identification of the TMEM95 egg receptor and may enable the development of infertility treatments and contraceptives for humans.

## Introduction

Fertilization is a central event of sexual reproduction, but how sperm and eggs bind to and fuse with one another has been largely undefined. Sperm IZUMO1 (1) and egg JUNO (2) mediate the only known cell surface interaction between mammalian gametes. Recent reports suggested that *Tmem95* (encoding transmembrane protein 95) mutant cattle exhibit impaired male fertility (3, 4); *Tmem95* knockout mice show male-specific sterility, producing sperm that can bind to, but do not fuse with eggs (5, 6). *Tmem95* encodes a sperm acrosomal membrane protein, which re-localizes to the equatorial segment of the sperm head (3, 6) where membrane fusion with the egg takes place (7, 8). These observations shed light on a potential role of TMEM95 in sperm-egg membrane fusion.

Humans also express *TMEM95* transcripts (9). In this study, we utilized the sperm penetration assay (10), a clinical laboratory test that evaluates fusion of human sperm with eggs from Syrian golden hamsters (*Mesocricetus auratus*), as a model system. TMEM95 is a type-I single-pass transmembrane protein (3, 5, 6). Motivated by a hypothesis that the ectodomain of TMEM95 binds to eggs through a specific, membrane-bound receptor on eggs, we found that a bivalent TMEM95 ectodomain protein binds hamster eggs, providing direct evidence for a TMEM95 receptor on eggs. The 1.5 Å-resolution X-ray crystal structure of TMEM95 we describe here reveals an evolutionarily conserved region of the protein with a positively charged surface. Amino-acid substitutions within this region of TMEM95 ablate egg binding. We speculate that this region serves as an egg-receptor binding site for TMEM95.

We also found that human TMEM95 plays a role in membrane fusion. After generating two monoclonal antibodies that bind to different epitopes of TMEM95, we observed that neither antibody blocks binding of human sperm to hamster eggs, but both could inhibit membrane fusion of sperm with eggs. Taken together, our results provide evidence for a specific, receptor-mediated interaction of human sperm TMEM95 with eggs and inform strategies for the identification of this receptor. We propose that the interaction of TMEM95 with eggs facilitates membrane fusion of human sperm and eggs.

## Results

### A bivalent TMEM95 protein binds hamster eggs

We hypothesized that the ectodomain of TMEM95 mediates a cell-surface interaction of sperm with eggs. To monitor the interaction between TMEM95 and eggs, we designed and produced TMEM95-Fc, a fusion protein of the ectodomain of human TMEM95 and the fragment crystallizable region of human immunoglobulin G1 (IgG1) (*SI Appendix*, Fig. S1A). TMEM95-Fc contains two copies of the TMEM95 ectodomain (Fig.1B) and the Fc confers increased avidity for binding over monomeric TMEM95. Given that human sperm can fuse with eggs from Syrian golden hamsters (10, 11), we incubated the Fc or TMEM95-Fc proteins with hamster eggs, whose surrounding zona pellucida and cumulus cells were removed. Using a fluorescently labeled anti-Fc antibody, we detected binding to the egg cell surface only with TMEM95-Fc, not Fc alone (Fig. 1A and B and *SI Appendix*, Fig. S1E). To confirm that our labeling approach can also detect known protein-protein interactions of sperm with eggs, we next surveyed IZUMO1-Fc on eggs, a fusion protein of sperm IZUMO1 (1) ectodomain with Fc. While IZUMO1-Fc binds eggs, the IZUMO1^W148A^-Fc variant does not (Fig. 1C and D). The substitution of W148A ablates the interaction of IZUMO1 with JUNO (*SI Appendix*, Fig. S1B-H) (12, 13), the egg receptor of IZUMO1 (2). Our results showed that TMEM95 binds egg plasma membranes and suggest the presence of a receptor for TMEM95 on eggs.

**Fig. 1.**
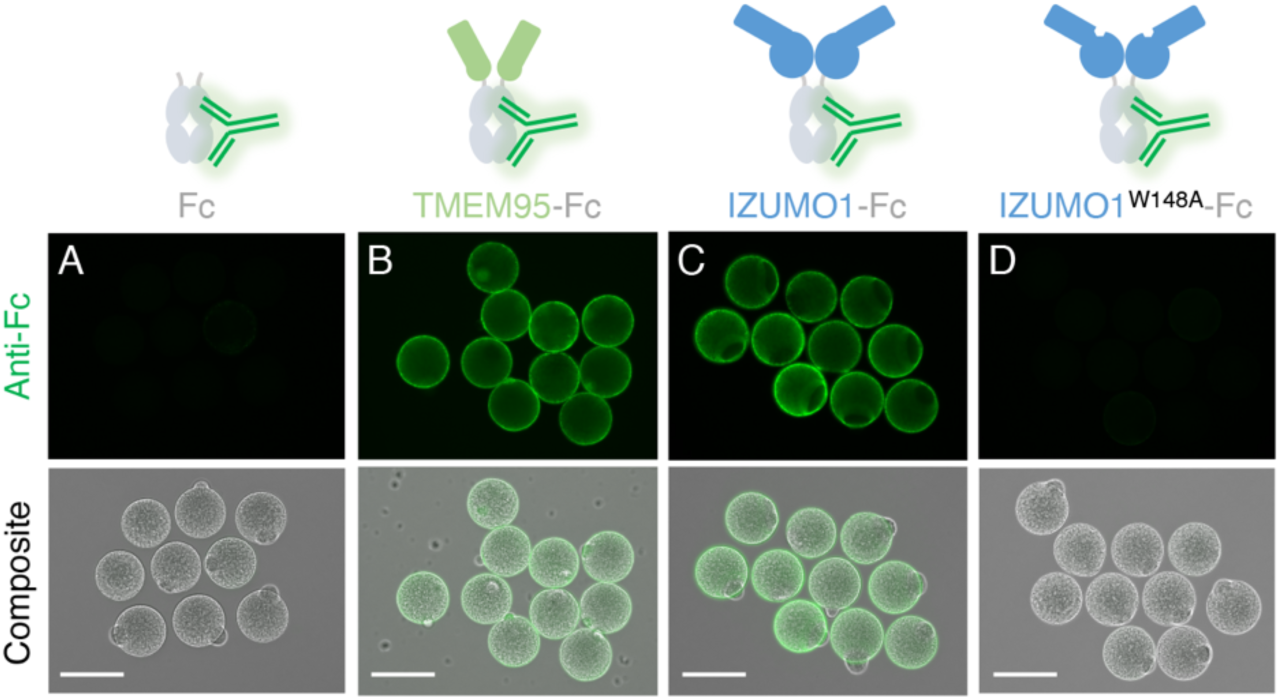
TMEM95-Fc binds eggs. Schematics of the Fc fusion protein with a fluorescence-conjugated Fc antibody. Immuno-fluorescence (upper) and differential interference contrast composite images (lower) of zona-free hamster eggs with 200 nM of (A) Fc, (B) TMEM95-Fc, (C) IZUMO1-Fc, or (D) IZUMO1^W148A^-Fc. Green fluorescence was conferred by a DyLight 488-conjugated Fc antibody. Scale bars, 100 μm. TMEM95-Fc and IZUMO1-Fc bind zona-free hamster eggs. See also *SI Appendix*, Fig. S1.

### The structure of TMEM95 is homologous to that of the N-terminus of IZUMO1

To understand how TMEM95 binds eggs, we determined a crystal structure of the TMEM95 ectodomain to 1.5 Å resolution using multi-wavelength anomalous X-ray diffraction (Fig. 2A and *SI Appendix*, Fig. S2A and Table S1). TMEM95 adopts an elongated rod shape, comprised of an N-terminal α-helical bundle (residues 17-110) and a C-terminal β-hairpin region (residues 111-135) (Fig. 2C). TMEM95 shows homology to the N-terminus of IZUMO1 (12, 13) with a C_α_ root-mean-square deviation of 7.2 Å, but TMEM95 does not share an immunoglobulin-like domain at the C-terminus with IZUMO1 (Fig. 2D). Unlike IZUMO1, the helical bundle of TMEM95 has three helices (α1, α3, and α4) and a coil (loop 2) that are arranged in an anti-parallel manner (α1-loop 2 and α3-α4). TMEM95 has three unique disulfide bonds: C35-C45 between α1 and loop 2 (*SI Appendix*, Fig. S2B), and C105-C134 and C109-C128 adjacent to the β-hairpin (Fig. 2B-D).

**Fig. 2.**
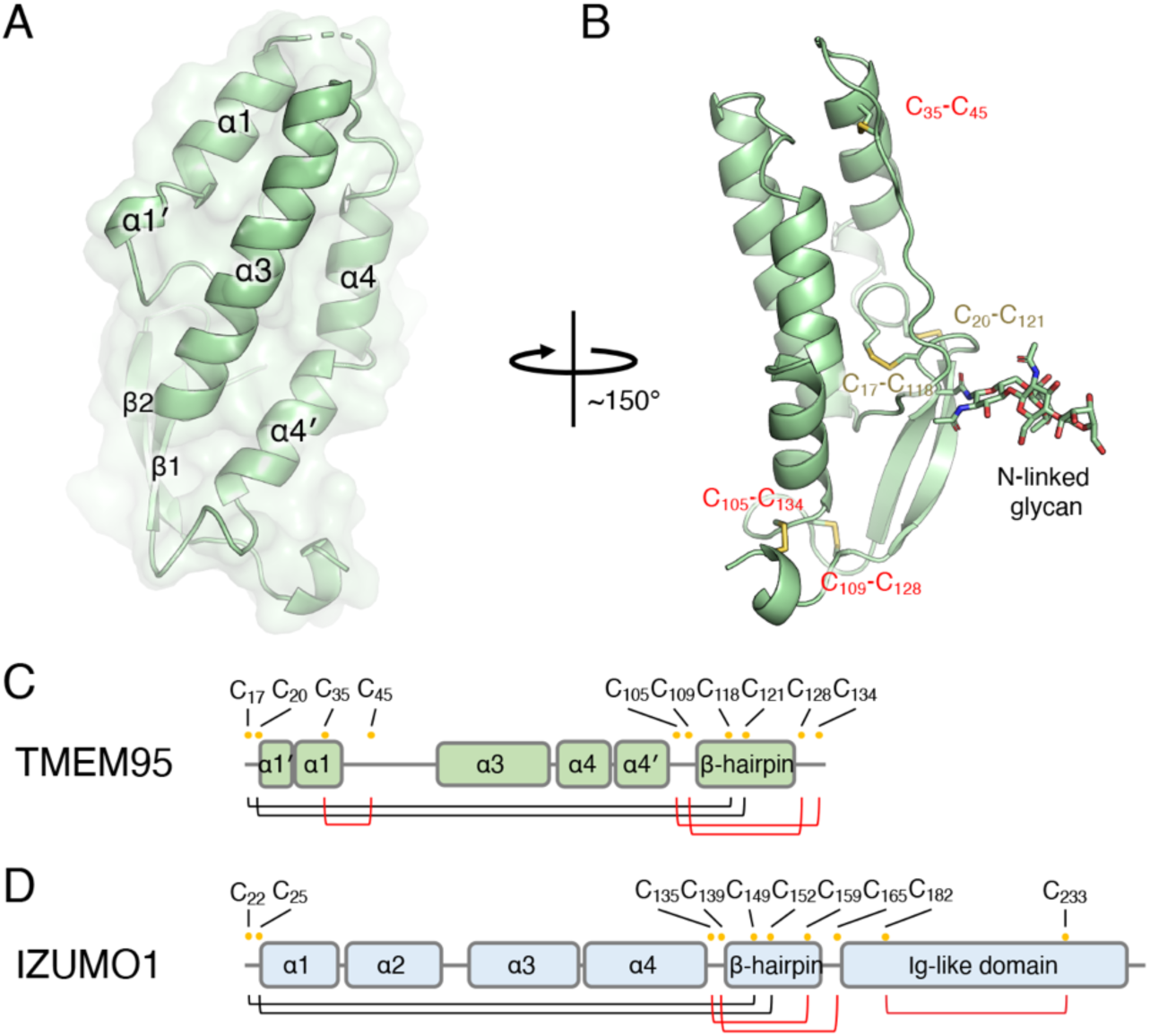
The structure of TMEM95 is homologous to IZUMO1. (A) Overlay of ribbon and space-filling diagrams of TMEM95 with structural elements labeled. (B) Ribbon diagram of TMEM95 with disulfide linkages labeled in yellow texts (same in IZUMO1) or red texts (different in IZUMO1). Domain organizations of (C) TMEM95 and (D) IZUMO1 with cysteine positions labeled as yellow dots and disulfide linked in black (same in TMEM95 and IZUMO1) or red lines (different in TMEM95 and IZUMO1). TMEM95 shows homology to the N-terminus of IZUMO1. See also *SI Appendix*, Fig. S2.

JUNO does not act as an egg receptor for TMEM95 (5, 6). A conserved N-linked glycan in the β-hairpin of TMEM95 (*SI Appendix*, Fig. S2E and F) could cause a clash if TMEM95 were to make a contact similar to that of IZUMO1 with JUNO (*SI Appendix*, Fig. S2C and D). However, even if this glycan is removed by the treatment with N-glycosidase PNGaseF, TMEM95-Fc does not bind JUNO (*SI Appendix*, Fig. S2G and H).

### A conserved surface of TMEM95 is a putative receptor-binding site

To gain further insights into the TMEM95 interaction with eggs, we analyzed the protein sequences of TMEM95 orthologs and mapped the degree of conservation for each amino acid onto the structure of TMEM95. We found that the area surrounding the N-glycan is variable (Fig. 3A), while the opposite side harbors a conserved (Fig. 3B), positively charged surface (Fig. 3C).

**Fig. 3.**
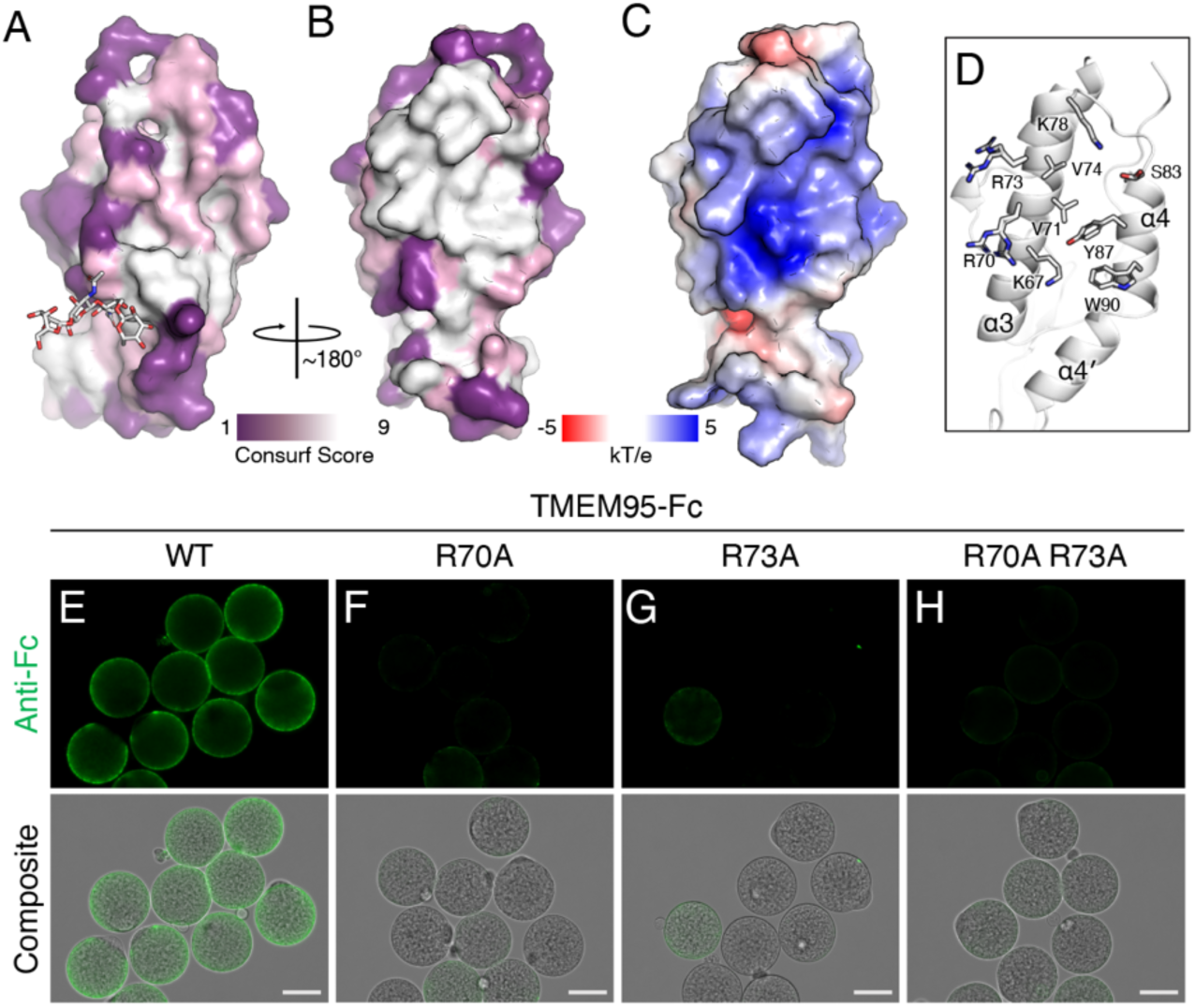
A conserved area of TMEM95 is a putative receptor binding site. (A-B) Space-filling *CONSURF* models (21) of TMEM95 with ∼180° rotation with purple representing variable and white representing conserved. (C) Space-filling model of electrostatic surface potential generated by APBS (Adaptive Poisson-Boltzmann Solver) with blue representing positively charged and red representing negatively charged. (D) Ribbon diagram of the conserved area of TMEM95 showing the side chains of surface-exposed residues. (E-H) Immuno-fluorescence (upper) and differential interference contrast composite images (lower) of zona-free hamster eggs with 200 nM of (E) TMEM95-Fc, (F) TMEM95^R70A^-Fc, (G) TMEM95^R73A^ -Fc, and (H) TMEM95^R70A^ ^R73A^-Fc. Green fluorescence by a DyLight 488-conjugated Fc antibody. Scale bars 50 μm. Substitutions of the conserved arginine residues on the identified surface of TMEM95 ablate egg-binding activities. See also *SI Appendix*, Fig. S3.

To examine whether the conserved, charged surface is critical for binding of TMEM95 to eggs, we produced TMEM95-Fc proteins that carry amino-acid substitutions of arginine residues (Fig. 3D and *SI Appendix*, Fig. S3A). These TMEM95 variants have melting temperatures comparable to that of the wild-type TMEM95-Fc protein (*SI Appendix*, Fig. S3B). When incubated with hamster eggs, the R70A, R73A, and R70A R73A TMEM95-Fc variants showed drastically reduced egg-binding activities compared to the wild-type (Fig. 3E-H and *SI Appendix*, Fig. S3C). Our data suggest that the identified evolutionarily conserved, positively charged surface of TMEM95 may function as a receptor-binding site.

### Monoclonal antibodies detect TMEM95 in human sperm

To generate reagents to investigate the functions of TMEM95 in human sperm, we immunized mice with the TMEM95 ectodomain (*SI Appendix*, Fig. S4A-C) and generated hybridoma cell lines that produce TMEM95 ectodomain-specific monoclonal antibodies, 3A01 and 6B08 (*SI Appendix*, Table S2). We used biolayer interferometry to assess the binding of the antibodies to TMEM95 (Fig. 4A) and found that 3A01 and 6B08 bind TMEM95 via two non-competing epitopes (Fig. 4B) with association constants of 1.4 nM and 1.3 nM, respectively (*SI Appendix*, Fig. S4D and E). The binding of either 3A01 or 6B08 to TMEM95-Fc does not inhibit its binding to the eggs (*SI Appendix*, Fig. S4G). 3A01 and 6B08 bind similarly to TMEM95-Fc and the R70A and R73A TMEM95-Fc variants (*SI Appendix*, Fig. S4H). These results suggest that the 3A01 and 6B08 antibodies against TMEM95 do not compete for binding of TMEM95 with its egg receptor.

**Fig. 4.**
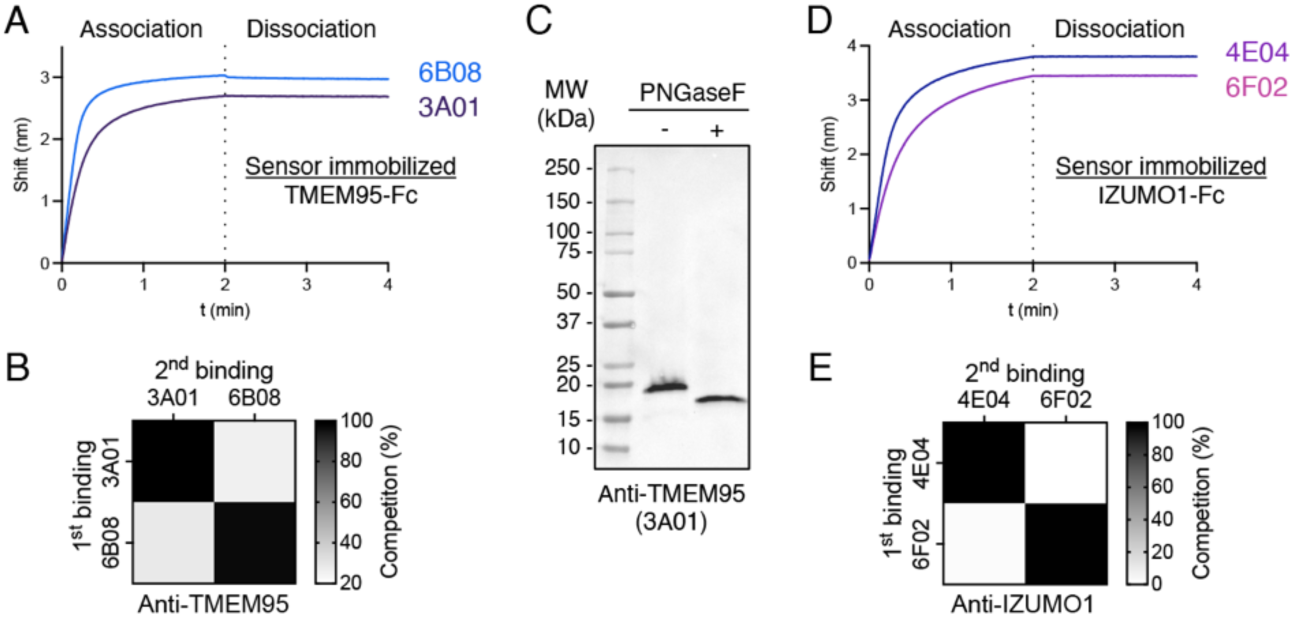
Antibodies detect the expression of TMEM95 in human sperm. (A, D) Biolayer interferometric traces of sensor immobilized (A) TMEM95-Fc binding to 200 nM of TMEM95 antibodies 3A01 IgG and 6B08 IgG or (D) IZUMO1-Fc binding to IZUMO1 antibodies 4E04 IgG and 6F02 IgG, with association for 2 min and dissociation for 2 min. (B, E) Summary in a heat map of antibody competition (B) of 3A01 IgG and 6B08 IgG to sensor immobilized TMEM95-Fc and (E) of 4E04 IgG and 6F02 IgG to sensor immobilized IZUMO1-Fc. (C) Human sperm lysates without or with PNGaseF treatments. Western blots were performed using non-heat-dentured, non-reduced sperm lysates by a primary antibody of 10 μg/mL anti-TMEM95 3A01 IgG, and a secondary HRP-conjugated anti-mouse antibody. TMEM95 is expressed and N-linked glycosylated in human sperm. See also *SI Appendix*, Figs. S4 and S5.

We next performed Western blotting using the TMEM95 antibodies to probe whole cell lysates of human sperm and each could detect a band of ∼20 kDa (*SI Appendix*, Fig. S4F), the expected molecular weight of TMEM95. To investigate whether TMEM95 is N-linked glycosylated, we treated the human sperm lysate with PNGaseF and observed a shift in size to ∼17.5 kDa (Fig. 4C), consistent with the loss of one glycan. Our results show that TMEM95 is expressed and N-linked glycosylated in human sperm.

Using a similar approach for IZUMO1 (*SI Appendix*, Fig. S5A-C), we generated hybridoma cell lines that produce IZUMO1-specific monoclonal antibodies, 4E04 and 6F02 (Fig. 4D and *SI Appendix*, Table S2). These antibodies both bind IZUMO1 (*SI Appendix*, Fig. S5F, S5J) via two non-competing epitopes (Fig. 4E and *SI Appendix*, Fig. S5D and E). Compared to 4E04-bound IZUMO1-Fc, 6F02-bound IZUMO1-Fc blocks binding of IZUMO1-Fc to eggs (*SI Appendix*, Fig. S5G) and JUNO (*SI Appendix*, Fig. S6H and I). These results suggest that 4E04 and 6F02 bind to different epitopes of IZUMO1, and that the 6F02 epitope overlaps with the IZUMO1-binding site for JUNO.

### TMEM95 antibodies impair fusion of human sperm to hamster eggs

To examine whether human TMEM95 plays a role in membrane fusion, we produced the fragments antigen-binding (Fab) of the TMEM95 and IZUMO1 antibodies and tested these in a sperm penetration assay. These Fab fragments bind antigens at nanomolar affinities (*SI Appendix*, Figs. S4E and S5E) and may have less steric effects in membrane fusion than their larger IgG counterparts. We inseminated hamster eggs with human sperm preincubated with the TMEM95 antibody Fab, 3A01 (Fig. 5C) or 6B08 (Fig. 5D). We used an untreated group as a negative control (Fig. 5A) and IZUMO1 antibody Fab 6F02-treatment as a positive control (Fig. 5B). Based on the numbers of bound (Fig. 5E) and fused (Fig. 5F) sperm per egg, we found that the TMEM95 antibody Fab fragments do not block binding of sperm to eggs (Fig. 5E).

**Fig. 5.**
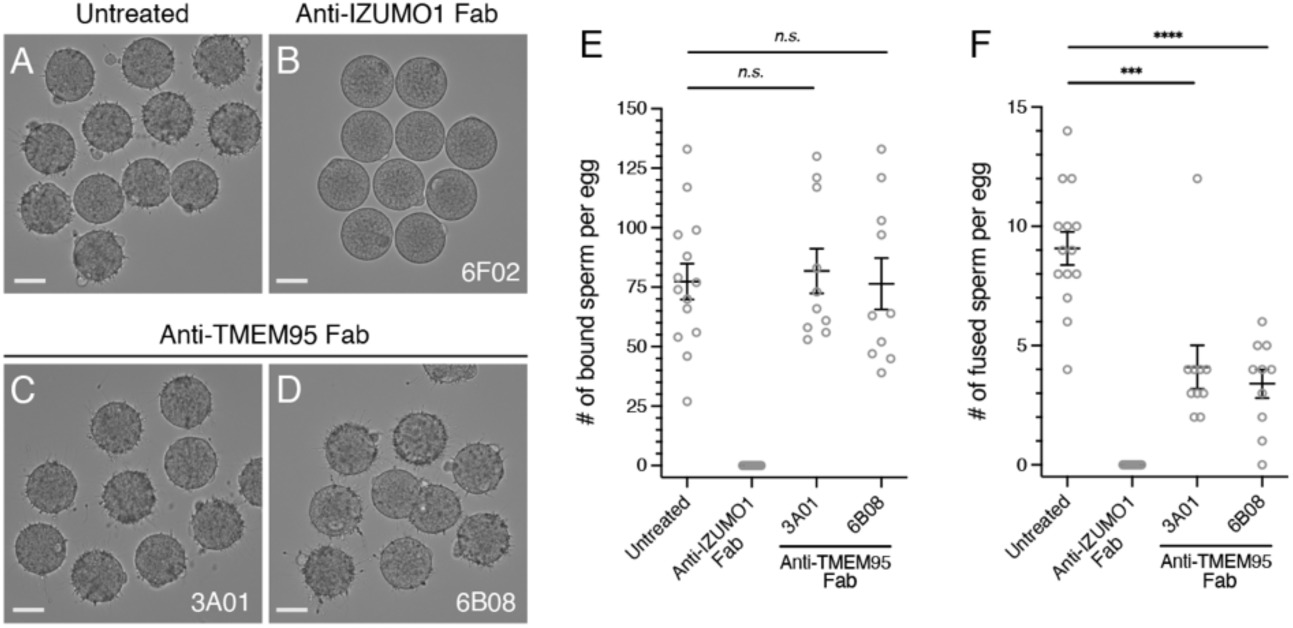
TMEM95 antibodies impair sperm-egg fusion. (A-D) Representative images showing binding of human sperm to zona-free hamster eggs (A) untreated or treated with 40 μg/mL of (B) anti-IZUMO1 Fab 6F02, (C) anti-TMEM95 Fab 3A01, or (D) anti-TMEM95 Fab 6B08. (E) Summary of the numbers of bound human sperm per zona-free hamster eggs (mean ± SEM), untreated 77.4 ± 7.5 (*N* = 14), anti-IZUMO1 Fab 6F02 0 ± 0 (*N* = 10), anti-TMEM95 3A01 Fab 81.8 ± 9.4 (*N* = 10, *n.s.*, not significant), and anti-TMEM95 6B08 Fab 76.4 ± 10.8 (*N* = 10, *n.s.*, not significant). (F) Summary of the numbers of fused human sperm per zona-free hamster eggs (mean ± SEM), untreated 9.1 ± 0.7 (*N* = 14), anti-IZUMO1 Fab 6F02 0 ± 0 (*N* = 10), anti-TMEM95 3A01 Fab 4.1 ± 0.9 (*N* = 10, *p* = 0.0002), and anti-TMEM95 6B08 Fab 3.4 ± 0.6 (*N* = 10, *p* < 0.0001). TMEM95 antibodies do not block sperm-egg binding but impair sperm-egg fusion. See also *SI Appendix*, Fig. S6.

However, the averaged numbers of fused sperm per egg significantly decreased from 9.1 ± 0.7 (mean ± standard error of the mean, SEM) in the untreated group to 4.1 ± 0.9 (*p* = 0.0002) and 3.4 ± 0.6 (*p* < 0.0001) in the TMEM95 Fab 3A01 and 6B08 groups, respectively (Fig. 5F and *SI Appendix*, Fig. S6A-D). Similarly, we observed that the TMEM95 antibody IgGs do not block sperm-egg binding (*SI Appendix*, Fig. S6E-G), but they decrease the average numbers of fused sperm per egg when compared with a control group treated with pre-immune IgG (*SI Appendix*, Fig. S6H-L). Therefore, the two non-competing TMEM95 monoclonal antibodies do not block sperm-egg binding but impair sperm-egg fusion, suggesting that TMEM95 plays a role in sperm-egg membrane fusion.

## Discussion

### Evidence for a receptor for TMEM95 on eggs

Our results provide compelling evidence for the existence of a membrane-bound receptor for sperm TMEM95 on eggs. Although the receptor has yet to be identified, our structural and site-directed mutagenesis studies identify a putative receptor-binding site on TMEM95. This region has a solvent-accessible surface area of ∼1,200 Å^2^, comparable to protein surfaces that mediate many protein-protein interactions (14, 15). We envision that the TMEM95 receptor is a membrane protein with a negatively charged region on its ectodomain surface. Nevertheless, we cannot rule out potential non-protein receptor candidates with electrostatic negative properties on the egg surface, such as phospholipids and glycans.

The bivalent TMEM95-Fc protein introduced here may be a useful reagent to facilitate the identification of the egg receptor of TMEM95. As cell surface interactions between membrane-bound proteins are often transient and dynamic (2, 16), the avidity of a bivalent protein could serve to stabilize the potentially weak interaction of TMEM95 with its receptor. TMEM95-Fc could therefore be used as a bait for the egg receptor, for example, for co-immunoprecipitation of mammalian eggs (*e.g.*, (17)), or for screening cultured cells expressing an egg cDNA library (*e.g.*, (2)).

### Potential roles of TMEM95 in membrane fusion

The TMEM95 antibodies used in this study do not ablate binding of TMEM95 to hamster eggs. How would the non-blocking antibodies of TMEM95 inhibit sperm-egg fusion? One possibility is that TMEM95 undergoes structural changes that are important for membrane fusion. Should sperm-egg fusion be accompanied by changes of TMEM95 in protein conformation or oligomeric state, the antibodies raised here against a defined conformation of TMEM95 may trap TMEM95 in a pre-fusion, monomeric state. Notably, early studies have suggested essential structural changes for IZUMO1 (*e.g.*, rearrangement of disulfides, protein dimerization) during sperm-egg membrane fusion (12, 18, 19).

Alternatively, or in addition, TMEM95 may assemble into a complex with other sperm proteins, such as a membrane fusogen. Antibody binding to TMEM95 could affect these events and explain the inhibitory results. Additionally, these antibodies might create steric hinderance which could interfere with membrane fusion (note, however, that an anti-IZUMO1 IgG, Mab125, does not block sperm-egg fusion, (20)).

Taken together, we conceptualize that sperm-egg membrane fusion involves pairwise cell surface interactions (Fig. 6). Sperm IZUMO1 binds egg JUNO, which mediates gamete adhesion, and a receptor-mediated interaction of sperm TMEM95 to the egg takes place; membrane fusion occurs thereafter. We anticipate additional analogous, yet to be identified, interactions between sperm proteins and their specific egg receptors.

**Fig. 6.**
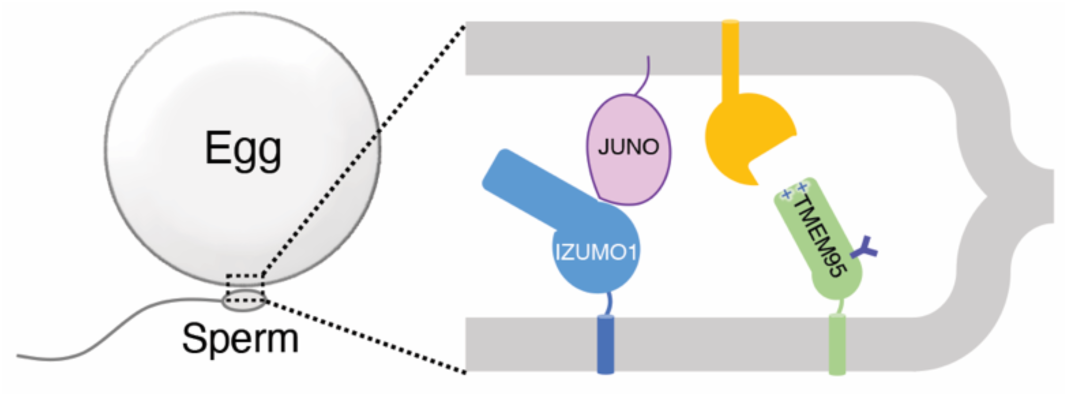
Model of sperm-egg binding and fusion. Illustration of membrane fusion of sperm and an egg and pairwise protein-protein interactions: sperm IZUMO1 (blue) binds egg JUNO (pink) and a receptor (orange)-mediated interaction of sperm TMEM95 (green) to the egg takes place; membrane fusion occurs thereafter.

In summary, our results suggest that human sperm TMEM95 likely plays a direct role in membrane fusion with eggs. Future work is needed to rule out indirect effects of TMEM95 antibodies that inhibit fusion while not blocking sperm-egg binding. More broadly, our work takes steps towards fully understanding the molecular interactions of the fertilization complex and has implications for the development of infertility treatments and contraceptives.

## Materials and methods

Additional information is provided in *SI Appendix, SI Materials and Methods*.

### Immunofluorescence microscopy of hamster eggs

Sexually mature female Syrian golden hamsters (Japan SLC Inc.) (approved by the Animal Care and Use Committee of the Research Institute for Microbial Diseases, Osaka University #28-4-2) were superovulated by peritoneal injection of pregnant mare serum gonadotropin and human coagulating gland (20 units for each; ASKA Pharmaceutical). Cumulus-oocyte complexes were extracted from the oviductal ampulla and treated with 1 mg/mL collagenase to remove the cumulus cells and zona pellucida, which yields zona-free eggs. These zona-free eggs were incubated with 200 nM Fc-fusion proteins in Biggers-Whitten-Whittingham (BWW) medium for 1 h and then stained with goat anti-human IgG Fc antibody DyLight 488 (Invitrogen) at a dilution of 1:50 for 1 h at 37 °C, 5% CO_2_. The eggs were imaged under a Keyence BZ-X810 microscope.

### Protein crystallization of TMEM95

Native TMEM95 proteins were crystallized at room temperature in a sitting-drop vapor diffusion system. 350 nL of 6.8 mg/mL protein was mixed with 350 uL of a reservoir solution of 150 mM NaCl, 20 mM HEPES pH 7.3, 30 mM CaCl_2_, 2% (w/v) PPG-P400, and 22% (w/v) PEG 3,350, over 80 μL of reservoir solution. Native crystals were supplemented with 20% (w/v) PEG 400 before cryo-cooling in liquid nitrogen. For multi-wavelength anomalous diffraction, crystals were grown in 150 mM NaCl, 20 mM HEPES pH 7.3, 10 mM CaCl_2_, 2% (w/v) PPG-P400, and 18% (w/v) PEG 3,350, and were transferred to a solution supplemented with 500 mM SmCl_3_ and incubated for ∼5 min. The Sm^3+^-bound crystals were washed in a SmCl_3_-free reservoir solution, and cryo-protected with 20% PEG 400 for cooling in liquid nitrogen.

### Sperm penetration assay

Sperm penetration assays were performed as previously described (10) with minor changes. Briefly, human semen from healthy donors, who had informed consent, was liquefied for 30 min at room temperature. The sperm were purified by discontinuous Percoll gradients (21) and incubated in BWW medium containing 2.5 μM calcium ionophore for 3 h at 37 °C, 5% CO_2_. The sperm were washed in fresh BWW medium and treated with monoclonal antibodies at 40 μg/mL for 1 h at 37 °C, 5% CO_2_. Zona-free hamster eggs were inseminated by the antibody-treated sperm at a density of 3×10^6^ motile sperm per mL for 3 h at 37 °C, 5% CO_2_. The eggs were washed in fresh BWW, gently flattened by coverslips, and examined under a phase-contrast microscope.

### Accession number

The coordinate and structure factor of human sperm TMEM95 ectodomain has been deposited in the RCSB Protein Data Bank under PDB ID code 7UX0. The structure is available immediately at https://peterkimlab.stanford.edu.

## Acknowledgement

We thank members of the P.S.K., M.I., and P.V.L. laboratories, Dr. Jonathan Z. Long, and Dr. Masaru Okabe for discussion, Dr. Mirella Bucci for comments on the manuscript, Gita Abhiraman and the laboratory of K. Christopher Garcia for protocols of baculovirus protein production, Dr. Daniel Fernandez of the Sarafan ChEM-H Macromolecular Structure Knowledge Center, and Silvia Russi of the Stanford Synchrotron Radiation Lightsource (SSRL) beam line 12-2 for X-ray crystallographic data collection. Use of the SSRL, SLAC National Accelerator Laboratory, is supported by the US Department of Energy (DOE), Office of Science, Office of Basic Energy Sciences under Contract DE-AC02-76SF00515. The SSRL Structural Molecular Biology Program is supported by the DOE, Office of Biological and Environmental Research and by a National Institutes of Health (NIH) grant P30GM133894.

We are grateful to the late Dr. Stuart Moss of the National Institute of Child Health and Human Development (NICHD). This work was supported by a NIH NICHD grant K99HD104924 (S.T.), Damon Runyon Cancer Research Foundation DRG-2301-17 (S.T.), the Ministry of Education, Culture, Sports, Science, and Technology, Japan Society for the Promotion of Science grants JP22K15103 (Y.L.), JP19H05750 (M.I.), and JP21H05033 (M.I.), a National Science Foundation Graduate Research Fellowship (W.M.S.), a Pew Biomedical Scholars Award (P.V.L.), the Global Consortium for Reproductive Longevity and Equality at the Buck Institute by the Bia-Echo Foundation (P.V.L.), the Virginia & D.K. Ludwig Fund for Cancer Research (P.S.K.), and Chan Zuckerberg Biohub (P.S.K.).

## SI Materials and Methods

### Expression and purification of the Fc-fusion proteins

The cDNAs encoding human TMEM95 (residues 1-145) or human IZUMO1 (residues 1-255) were subcloned into a pADD2 vector that carries a C-terminal fusion of a TEV protease cleavage site, a human IgG1 Fc, an Avi tag, and a hexa-histidine tag (*SI Appendix*, Table S3). The recombinant Fc fusion proteins were overexpressed by transient transfection of HEK293F cells (ThermoFisher) cultured at 37 °C, 8% CO_2_. TMEM95-Fc was purified by Ni-NTA affinity purification (Invitrogen), followed by anion exchange using an AKTA pure system by a Mono Q 5/50 GL (Cytiva). IZUMO1-Fc was purified by Protein-A affinity purification using a MabSelect Prism (Cytiva), followed by anion exchange using a Mono Q 5/50 GL. Purified proteins were stored in a buffer of 150 mM NaCl, 20 mM HEPES pH 7.4.

Tagless IZUMO1 proteins were obtained from IZUMO1-Fc through TEV (Sigma-Aldrich) cleavage overnight at 4 °C. Undigested proteins and the histidine tagged TEV proteases were removed by a MabSelect Prism followed by a HisTrap excel (Cytiva). Tagless IZUMO1 was further purified by gel filtration with a Superdex 200 Increase 10/300 GL (Cytiva) in a buffer of 150 mM NaCl, 20 mM HEPES pH 7.4.

### Protein expression and purification of JUNO

The cDNA encoding human JUNO (residues 20-227) was subcloned into a baculoviral vector pACgp67a that carries a signal sequence of MVSAIVLYVLLAAAAHSAFA and C-terminal hexa-histidine tag (*SI Appendix*, Table S3). Baculovirus was generated from Sf9 cells (ThermoFisher) by a co-transfection of pACgp67a and the BestBac Linearized Baculovirus DNA (Expression Systems). Passage one baculovirus was tittered and used for infecting HighFive cells (ThermoFisher) cultured at 27 °C. ∼3 days post infection, the conditioned media were harvested and mixed with NiCl_2_, CaCl_2_, and Tris pH 8.0 to a final concentration of 1 mM, 5 mM, and 100 mM, respectively. After centrifugation, the JUNO-His_6_ proteins was purified by Ni-NTA affinity purification from the resulting supernatant, followed by gel filtration with a Superdex 200 Increase 10/300 GL in a buffer of 150 mM NaCl, 20 mM HEPES pH 7.4.

### Biolayer interferometry

An Octet RED96 system (Pall ForteBio) was employed for protein-protein interaction assays in a buffer of 150 mM NaCl, 20 mM HEPES pH 7.4, 0.1% bovine serum albumin, and 0.05% Tween 20 at 29 °C under a shaking speed of 1,000 rpm. Biotinylated TMEM95-Fc or IZUMO1-Fc proteins were loaded onto Streptavidin biosensors (Sartorius). After loading the biosensors were baselined, associated in defined concentrations of analytes, and dissociated in the buffer with no analytes. Baseline-corrected binding traces were plotted and analyzed using GraphPad Prism 9.

### Differential scanning fluorimetry

A Prometheus NT.48 (NanoTemper) was employed for nanoscale differential scanning fluorimetry (NanoDSF). Protein samples were loaded into capillaries and subject to a temperature from 20 to 95 °C at a heating rate of 1 °C/min. Intrinsic fluorescence at 350 nm and 330 nm was recorded as a function of temperature. Thermal melting profiles were plotted using the first derivative of the ratio (*F*_350 nm_/*F*_330 nm_). Melting temperatures were calculated by the instrument and represented peaks in the thermal melting curves.

### Protein purification of TMEM95

The cDNA encoding human TMEM95 (residues 17-138) was subcloned into a pADD2 vector. The N-terminus of TMEM95 was fused to a signal sequence of MRMQLLLLIALSLALVTNS and the C-terminus to a C-tag of EPEA (*SI Appendix*, Table S3). Recombinant TMEM95 proteins were overexpressed in HEK293F cells by transient transfection. Affinity purification was performed using the CaptureSelect C-tagXL affinity matrix (ThermoFisher). The eluate was purified by cation exchange by a Mono S 5/50 GL (Cytiva), followed by gel filtration with a Superdex 200 Increase 10/300 GL in a buffer of 150 mM NaCl, 20 mM HEPES pH 7.4. Size exclusion with multiangle light scattering was performed on an Agilent 1260 Infinity II high performance liquid chromatography coupled with Wyatt detectors for light scattering (miniDAWN) and refractive index (Optilab).

### X-ray crystallography

X-ray diffraction data were collected at the Stanford Synchrotron Radiation Lightsource (SSRL) beam line 12-2 of SLAC National Accelerator Laboratory. For the Sm^3+^-bound crystal, multi-wavelength anomalous diffraction data were collected at wavelengths 1.694 Å (peak), 1.137 Å (remote), and 1.695 Å (inflection). For the native crystal, the diffraction data were collected at 0.979 Å wavelength to 1.50 Å resolution. All diffraction data were processed using *autoPROC* (1). The TMEM95 ectodomain structure was solved by experimental phasing using *AutoSol* in *Phenix* (2). An initial model containing 110 amino acids and two Sm^3+^ ions were obtained using *AutoBuild* and was subsequently applied to the native X-ray dataset by molecular replacement using *Phaser*. Model refinement and density modification were performed in *Phenix*. Model building was performed using *Coot* (3). Structural illustrations were generated with *PyMOL*.

### Evolutionary conservation by *CONSURF*

The protein sequence of human TMEM95 was input as a query sequence for a protein BLAST search using blastp. The top 150 results from were filtered manually and ortholog-unique sequences were subjected for alignment by MAFFT. The multiple sequence alignment and the TMEM95 structure were used as input in the *CONSURF* server (4). The overall conservation scores from 1 to 9 were calculated using the Bayesian methods for each amino acid and were mapped onto the TMEM95 structure in a color-coordinated fashion as shown in Fig. 3.

### Generation of mouse hybridomas

Five female BALB/c mice (Jackson Laboratory) aged ∼8 weeks (approved by Stanford University Administrative Panel on Laboratory Animal Care, APLAC 33984) were immunized with 10 μg purified protein of TMEM95 (residues 17-138) in 100 μL of 150 mM NaCl, 20 mM HEPES pH 7.4, adjuvanted with 10 μg Quil-A (Invivogen) and 10 μg monophosphoryl lipid A (InvivoGen). Mice were boosted at days 21, 43, 64, and 86. At day 90, a spleen of one mouse was disaggregated into a single-cell suspension for hybridoma generation following the manufacturer’s procedures (Stemcell technologies). Briefly, splenocytes were purified and fused with Sp2/0-Ag14 cells (ATCC) using polyethylene glycol. Hybridomas were cultured in 96-well plates with a selection medium containing hypoxanthine, aminopterin, and thymidine. ∼14 days after recovery, the conditioned media were screened for binding to TMEM95 by ELISA. TMEM95-binding-positive cells were sorted as single cells in 96-well plates using a SONY SH800S. ∼14 days after recovery, the conditioned media were screened, and the selected TMEM95-positive clones were expanded for antibody sequencing (Genscript Biotech, Table S2). Similarly, five mice were immunized with IZUMO1 (residues 22-255) and boosted at days 21, 43, 61, and 96. At day 100, a spleen from one mouse was used for hybridoma generation.

### Antibody production and purification

Hybridomas producing the TMEM95 and IZUMO1 antibodies were cultured in ClonaCell-HY Medium E (Stemcell technologies) and subsequently adapted to serum-free AOF Expansion Medium (Stemcell technologies) for 5-7 days at 37 °C, 5% CO_2_. The IgG in the conditioned AOF media was harvested from the supernatants and subjected for affinity purification by a HiTrap Protein G HP (Cytiva) and gel filtration with a Superdex 200 Increase 10/300 GL in a buffer of 150 mM NaCl, 20 mM HEPES pH 7.4.

The cDNAs encoding the heavy and light chains of the TMEM95 and IZUMO1 antibody Fab were subcloned into a pVRC vector (*SI Appendix*, Table S3). The Fabs were produced in HEK293F cells by transient transfection at 37 °C, 8% CO_2_, and purified from the supernatants of the conditioned media by a HiTrap Protein G HP, followed by gel filtration with a Superdex 200 Increase 10/300 GL in a buffer of 150 mM NaCl, 20 mM HEPES pH 7.4. All antibodies were concentrated to 1.0 mg/mL, supplemented with 10% glycerol, and aliquoted for long-term storage at -80 °C.

### Human sperm isolation and western blotting

The experimental procedures utilizing human derived samples in Fig. 4 and *SI Appendix* Figs. S4 and S5 were approved by the Committee on Human Research at the University of California, Berkeley, IRB protocol 2013-06-5395. Purified human sperm (5) were lysed in a buffer of 150 mM NaCl, 50 mM Tris pH 7.4, 1% Triton X-100, 0.5% Sodium deoxylcholate, 0.1% SDS, 1 mM EDTA, 10% (v/v) glycerol, and Halt protease inhibitors (ThermoFisher). The protein concentrations of the whole cell lysates were estimated by a Bradford assay (BioRad) using bovine serum albumin as a standard. The lysates were stored at 4 °C in a non-reducing condition before loaded onto an SDS-PAGE gel for electrophoresis. 15 μg of lysates and 10 μg/mL of TMEM95 antibodies were used for the detection of TMEM95; 7 μg of lysates and 2 μg/mL of IZUMO1 antibodies were used for the detection of IZUMO1. A secondary antibody of HRP-conjugated goat anti-mouse IgG (BioLegend) was used for immunoblotting. PNGaseF treatment was performed under non-reducing conditions following the manufacturer’s instructions (NEB).

---

**Fig. S1.**
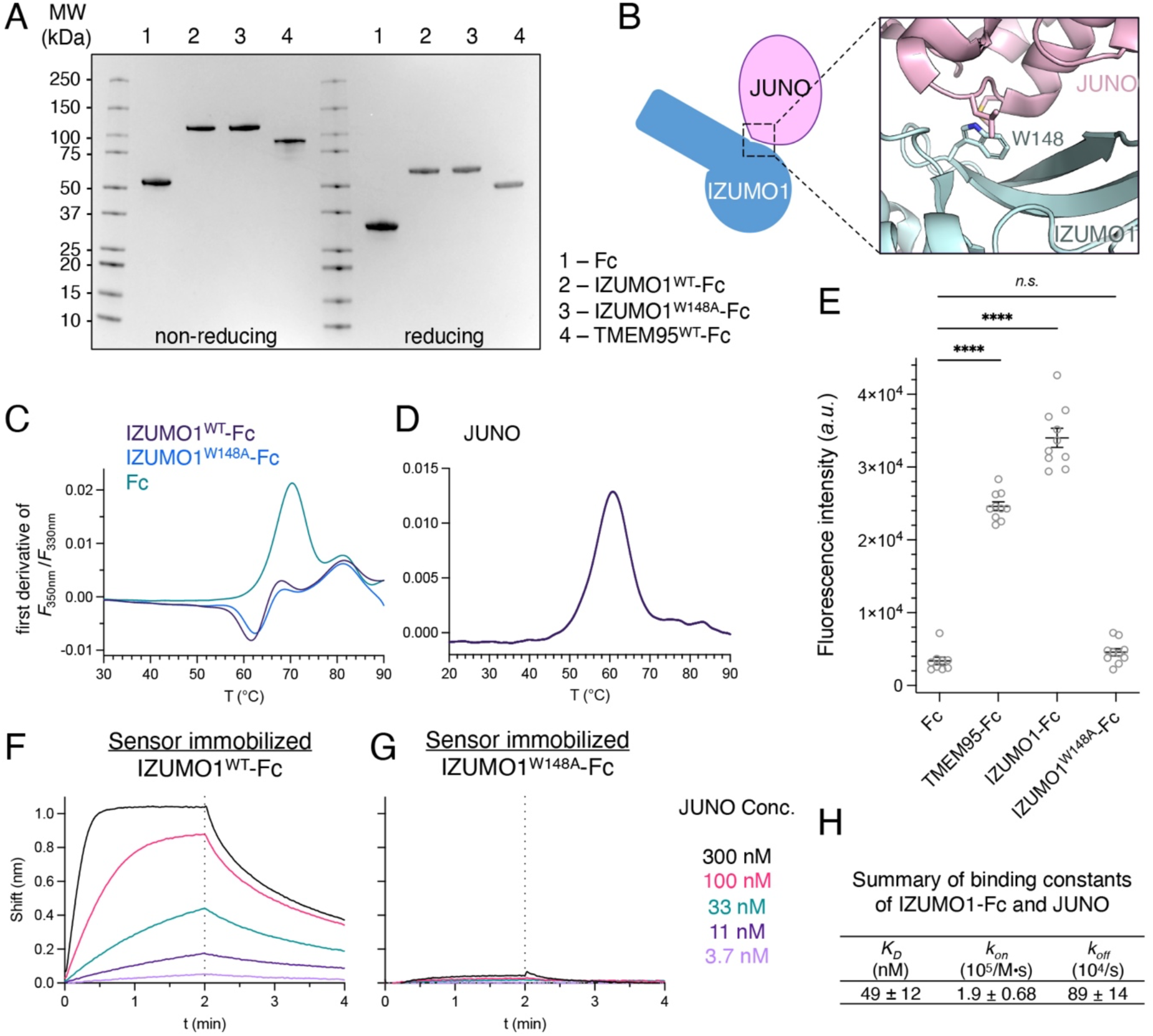
related to Fig. 1, Characterization of IZUMO1-Fc. (A) Coomassie-blue stained SDS-PAGE gel of Fc, IZUMO1^WT^-Fc, IZUMO1^W148A^-Fc, and TMEM95-Fc proteins under non-reducing (left) or reducing conditions (right). (B) Cartoon schematic and ribbon diagram (PDB ID: 5F4E) of the IZUMO1-JUNO complex showing the side chain of W148 of IZUMO1 mediates interactions with JUNO. (C-D) NanoDSF thermal melting profiles of (C) Fc, IZUMO1^WT^-Fc, IZUMO1^W148A^-Fc, and (D) JUNO proteins. (E) Quantification of fluorescence intensities (*a.u*., arbitrary unit; ****, *p* < 0.0001, *n.s.* not significant) of Fc, TMEM95-Fc, IZUMO1^WT^-Fc, and IZUMO1^W148A^-Fc on eggs shown in Figure 1. (F-G) Biolayer interferometric traces of sensor immobilized (F) IZUMO1-Fc or (G) IZUMO1^W148A^-Fc binding JUNO of 300 nM, 100 nM, 33 nM, 11 nM, and 3.7 nM, with association for 2 min and dissociation for 2 min. (H) List of binding constants of IZUMO1-Fc with JUNO calculated from traces in (F).

**Fig. S2.**
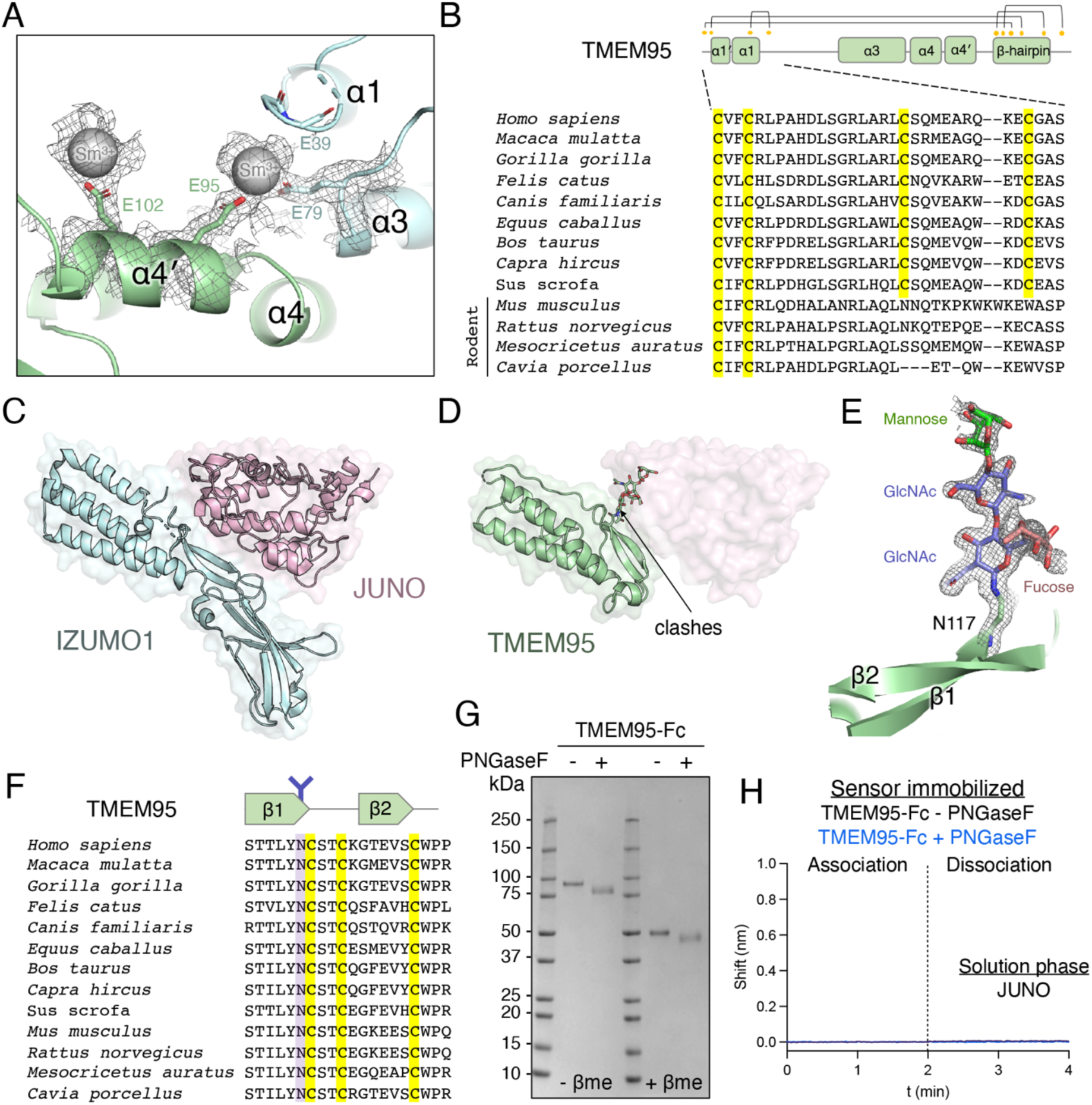
related to Fig. 2, JUNO is not a receptor for TMEM95. (A) Ribbon diagram overlay with a 2*F*_obs_-*F*_calc_ electron density map surrounding the Sm^3+^ ions between two TMEM95 protomers (green and cyan) in the crystal lattice solved by multi-wavelength X-ray anomalous diffraction. (B) Multiple sequence alignment of the α1 region of TMEM95 orthologs with conserved cysteines highlighted in yellow. (C-D) Ribbon diagrams overlay with a space-filling model of (C) the IZUMO1-JUNO complex (PDB ID: 5F4E), (D) the TMEM95-superimposed JUNO complex, where the N-glycan of TMEM95 causes a clash with JUNO. (E) Ribbon diagram overlay with a 2*F*_obs_-*F*_calc_ composite omit (10%) electron density map of the N117 side chain and its linked glycan. (F) Multiple sequence alignment of the β-hairpin of TMEM95 orthologs with conserved cysteines highlighted in yellow and the asparagine in purple. (G) Coomassie-blue stained SDS-PAGE gel of TMEM95-Fc treated without or with PNGaseF under non-reducing (left) or reducing (right) conditions. (H) Biolayer interferometric traces of sensor immobilized TMEM95-Fc treated without or with PNGaseF binding to JUNO of 300 nM, with association for 2 min and dissociation for 2 min.

**Fig. S3.**
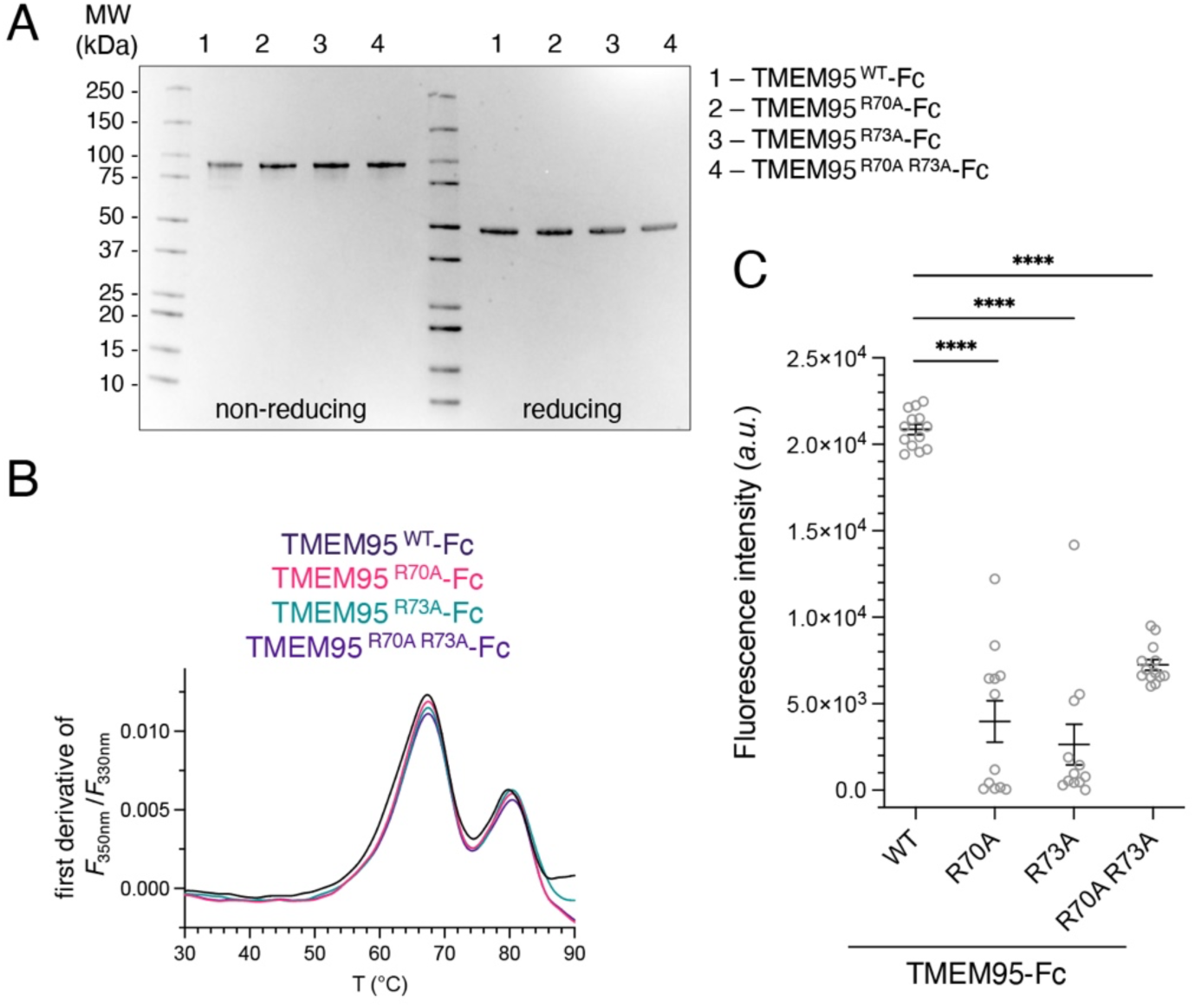
related to Fig. 3, Characterization of TMEM95-Fc. (A) Coomassie-blue stained SDS-PAGE gel of TMEM95-Fc, TMEM95^R70A^-Fc, TMEM95^R73A^-Fc, and TMEM95^R70A R73A^-Fc proteins under non-reducing (left) or reducing conditions (right). (B) NanoDSF thermal melting profiles of TMEM95-Fc, TMEM95^R70A^-Fc, TMEM95^R73A^-Fc, and TMEM95^R70A^ ^R73A^-Fc proteins. (C) Quantified green fluorescence intensities (*a.u*., arbitrary unit; ****, *p* < 0.0001) of TMEM95-Fc, TMEM95^R70A^-Fc, TMEM95^R73A^-Fc, and TMEM95^R70A^ ^R73A^-Fc proteins on eggs shown in Figure 3.

**Fig. S4.**
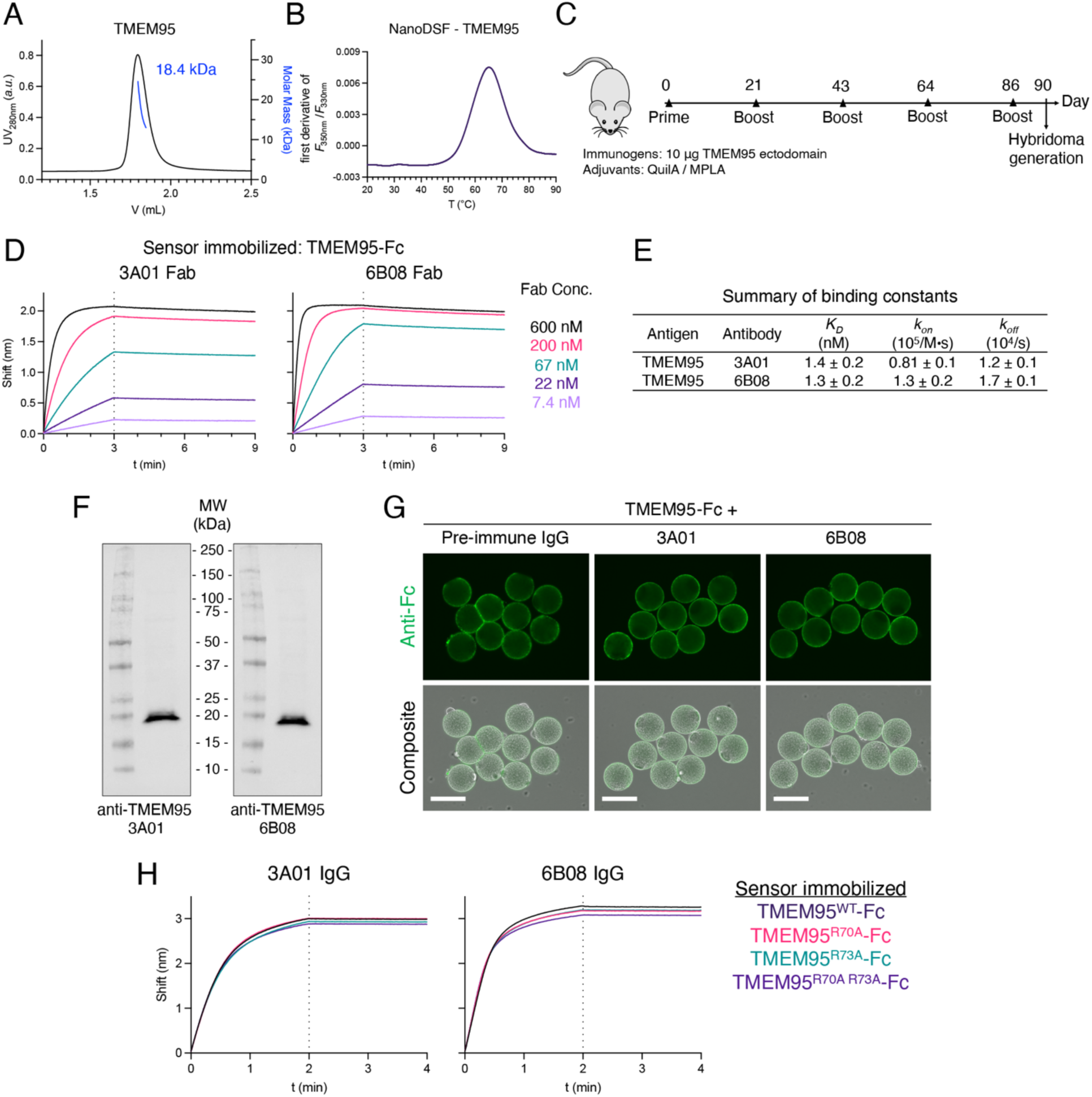
related to Fig. 4, Characterization of the TMEM95 antibodies. (A) Size exclusion and multiangle light scattering of the TMEM95 protein showing a monodispersed peak with a calculated molecular weight of 18.4 kDa, as an expected monomer in solution. (B) NanoDSF thermal melting profile of the TMEM95 protein used for protein crystallization and mouse immunization. (C) Schedule of mouse immunization using the TMEM95 protein. (D) Biolayer interferometric traces of sensor immobilized TMEM95-Fc binding to 3A01 Fab or 6B08 Fab at concentrations of 600 nM, 200 nM, 67 nM, 22 nM, and 7.4 nM, with association for 3 min and dissociation for 6 min. (E) Summary of binding constants of TMEM95-Fc with anti-TMEM95 3A01 Fab and 6B08 Fab calculated from traces in (D). (F) Western blots of non-heat-denatured, non-reduced human sperm lysates by a primary antibody of 10 μg/mL anti-TMEM95 3A01 IgG or 6B08 IgG, and a secondary HRP-conjugated anti-mouse antibody. (G) Immuno-fluorescence (upper) and differential interference contrast composite images (lower) of zona-free hamster eggs incubated with TMEM95-Fc that has been pre-bound to protein G purified pre-immune mouse IgG, anti-TMEM95 3A01 IgG, or 6B08 IgG. 2.5 μM TMEM95-Fc was mixed with 5 μM (0.75 mg/mL) IgG for 1 hour to form a complex of TMEM95-Fc and the antibody, and the mixture was added to the eggs at a final concentration of 200 nM TMEM95-Fc. Green fluorescence by a DyLight 488-conjugated Fc antibody. Scale bars 100 μm. (H) Biolayer interferometric traces of sensor-immobilized TMEM95-Fc, TMEM95^R70A^-Fc, TMEM95^R73A^-Fc, and TMEM95^R70A^ ^R73A^-Fc proteins binding to 200 nM of 3A01 IgG or 6B08 IgG, with association for 2 min and dissociation for 2 min.

**Fig. S5.**
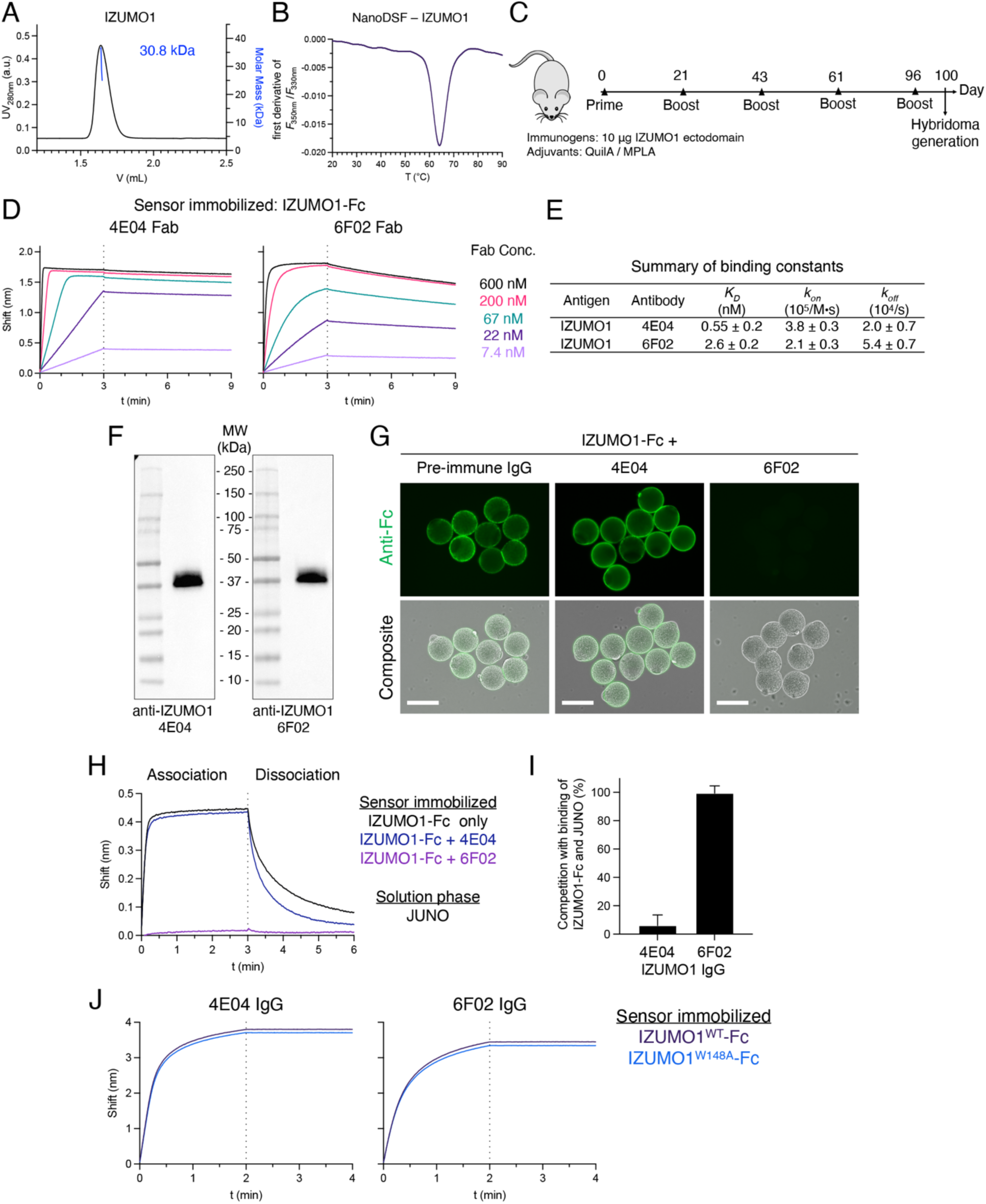
related to Fig. 4, Characterization of the IZUMO1 antibodies. (A) Size exclusion and multiangle light scattering of the IZUMO1 protein showing a monodispersed peak with a calculated molecular weight of 30.8 kDa, as an expected monomer in solution. (B) NanoDSF thermal melting profile of the IZUMO1 protein used for mouse immunization. (C) Schedule of mouse immunization of the IZUMO1 protein. (D) Biolayer interferometric traces of sensor immobilized IZUMO1-Fc binding to 4E04 Fab or 6F02 Fab at concentrations of 600 nM, 200 nM, 67 nM, 22 nM, and 7.4 nM, with association for 3 min and dissociation for 6 min. (E) Summary of binding constants of IZUMO1-Fc with anti-IZUMO1 4E04 Fab and 6F02 Fab calculated from traces in (D). (F) Western blots of non-heat-denatured, non-reduced human sperm lysates by a primary antibody of 2 μg/mL anti-IZUMO1 4E04 IgG or 6F02 IgG, and a secondary HRP-conjugated anti-mouse antibody. (G) Immuno-fluorescence (upper) and differential interference contrast composite images (lower) of zona-free hamster eggs incubated with IZUMO1-Fc that has been pre-bound to protein G purified pre-immune mouse IgG, anti-IZUMO1 4E04 IgG, or 6F02 IgG. 2.5 μM IZUMO1-Fc was mixed with 5 μM (0.75 mg/mL) IgG for 1 hour to form a complex of IZUMO1-Fc and the antibody, and the mixture was added to the eggs at a final concentration of 200 nM IZUMO1-Fc. Green fluorescence by a DyLight 488-conjugated Fc antibody. Scale bars, 100 μm. (H) Biolayer interferometric traces of sensor immobilized IZUMO1-Fc, or IZUMO1-Fc in complex with 4E04 IgG, 6F02 IgG binding to 300 nM JUNO, with association for 3 min and dissociation for 3 min. (I) Summary of antibody competition with the IZUMO1-Fc and JUNO interaction calculated from (H). (J) Biolayer interferometric traces of sensor-immobilized IZUMO1-Fc and IZUMO1^W148A^-Fc binding to 200 nM of 4E04 IgG or 6F02 IgG, with association for 2 min and dissociation for 2 min.

**Fig. S6.**
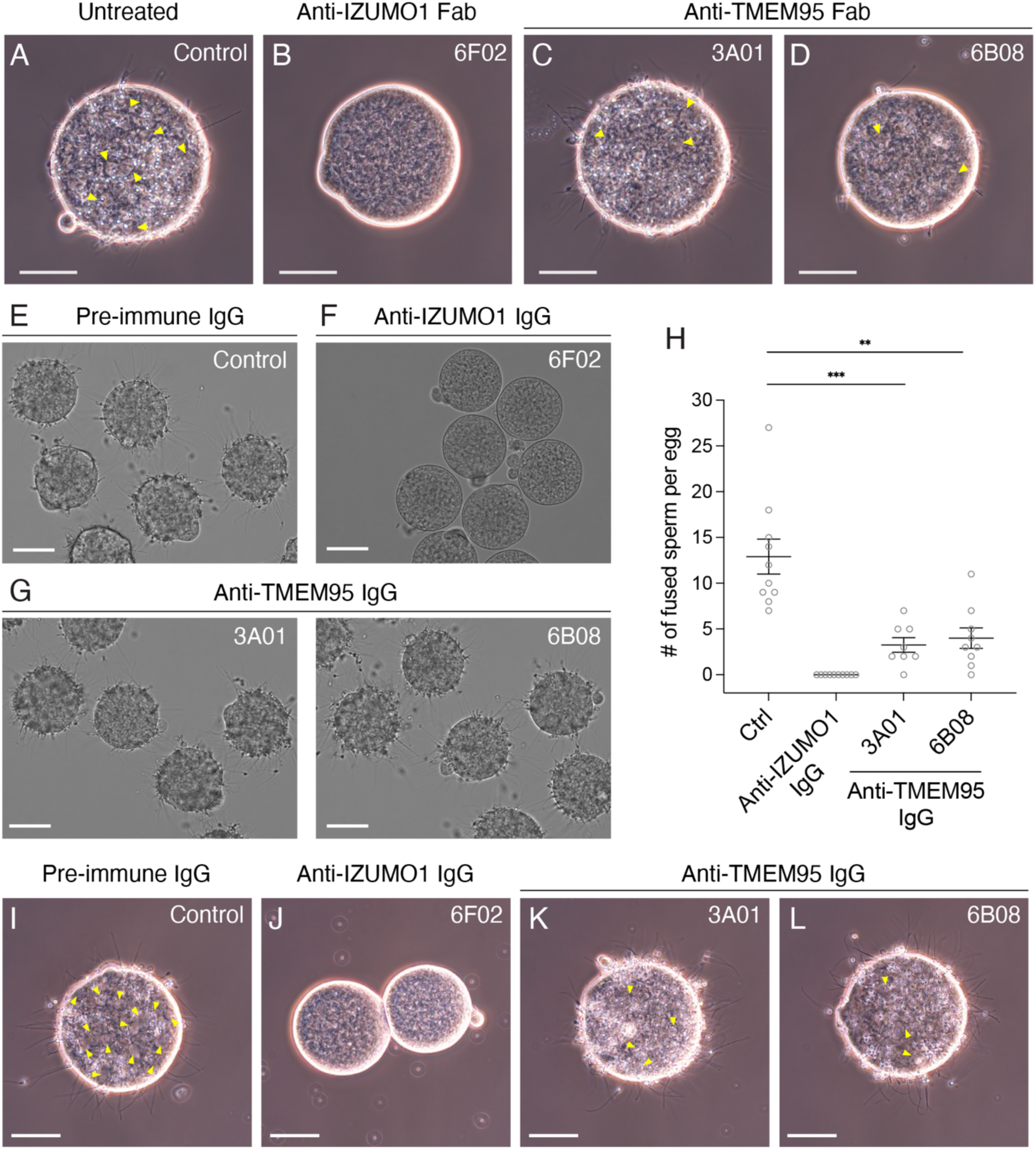
related to Fig. 5, TMEM95 antibodies impair sperm-egg fusion. Representative images showing fusion of human sperm with zona-free hamster eggs (A) untreated or treated with 40 μg/mL of (B) anti-IZUMO1 Fab 6F02, (C) anti-TMEM95 Fab 3A01, or (D) anti-TMEM95 Fab 6B08. Arrows indicating fused sperm with swollen sperm heads. Scale bars, 50 μm. (E-G) Representative images showing binding of human sperm with zona-free hamster eggs in the presence of 40 μg/mL (E) pre-immune mouse IgG, (F) anti-IZUMO1 IgG 6F02, (G) anti-TMEM95 IgG 3A01 (left), or anti-TMEM95 IgG 6B08 (right) after 3 hours of insemination. Scale bars, 50 μm. (H) Summary of the numbers of fused human sperm per zona-free hamster eggs in each group (mean ± SEM), control 12.9 ± 1.7 (*N* = 10), anti-IZUMO1 6F02 0 ± 0 (*N* = 10), anti-TMEM95 3A01 3.3 ± 0.8 (*N* = 8, *p* < 0.001), and anti-TMEM95 6B08 4.0 ± 1.1 (*N* = 9, *p* < 0.01). Representative images showing fusion of human sperm with zona-free hamster eggs in the presence of 40 μg/mL (I) pre-immune mouse IgG, (J) anti-IZUMO1 IgG 6F02, (K) anti-TMEM95 IgG 3A01, and (L) anti-TMEM95 IgG 6B08. Arrows indicating fused sperm with swollen sperm heads. Scale bars, 50 μm.

**Table S1.** Crystallographic data collection and refinement statistics

|  | Human TMEM95 ectodomain |  |  |  |
| --- | --- | --- | --- | --- |
|  | Native | Multi-wavelength anomalous diffraction |  |  |
|  |  | Peak | Remote | Inflection |
| **PDB ID** | 7UX0 |  |  |  |
| **Wavelength, Å** | 0.97946 | 1.69457 | 1.13743 | 1.69515 |
| **Resolution range, Å** | 36.01 - 1.50<br>(1.52 - 1.50) | 39.01 - 2.10<br>(2.14 - 2.10) | 29.09 - 1.97<br>(2.00 - 1.97) | 39.02 - 2.12<br>(2.15 - 2.12) |
| **Space group** | *P* 2<sub>1</sub> 2<sub>1</sub> 2<sub>1</sub> | *P* 2<sub>1</sub> 2<sub>1</sub> 2<sub>1</sub> | *P* 2<sub>1</sub> 2<sub>1</sub> 2<sub>1</sub> | *P* 2<sub>1</sub> 2<sub>1</sub> 2<sub>1</sub> |
| **Unit cell** | 39.53 43.11 72.02<br>90 90 90 | 39.30 43.22 78.01<br>90 90 90 | 39.32 43.25 78.09<br>90 90 90 | 39.31 43.23 78.03<br>90 90 90 |
| **Total reflections** | 258299 (13130) | 90875 (3275) | 109780 (6269) | 92437 (3364) |
| **Unique reflections** | 20505 (1004) | 7810 (370) | 8952 (476) | 7866 (365) |
| **Multiplicity** | 12.6 (13.1) | 11.6 (8.9) | 12.3 (13.2) | 11.8 (9.2) |
| **Completeness, %** | 99.7 (100.0) | 95.7 (92.7) | 90.1 (100.0) | 98.0 (93.8) |
| **Mean *I*/σ(*I*)** | 12.3 (2.2) | 14.20 (2.4) | 12.10 (2.2) | 14.1 (2.1) |
| ***R*<sub>merge</sub>** | 0.148 (1.803) | 0.188 (0.922) | 0.184 (1.641) | 0.173 (1.137) |
| ***CC*<sub>1/2</sub>** | 0.997 (0.855) | 0.998 (0.882) | 0.998 (0.834) | 0.998 (0.786) |
| ***R*<sub>work</sub>** | 0.223 |  |  |  |
| ***R*<sub>free</sub>** | 0.251 |  |  |  |
| **Number of non-hydrogen atoms** | 1133 |  |  |  |
| macromolecules | 985 |  |  |  |
| solvent | 89 |  |  |  |
| **Protein residues** | 115 |  |  |  |
| **RMS(bonds), Å** | 0.007 |  |  |  |
| **RMS(angles), °** | 1.03 |  |  |  |
| **Ramachandran favored, %** | 98.20 |  |  |  |
| **Ramachandran outliers, %** | 0.00 |  |  |  |
| **Clashscore** | 3.90 |  |  |  |
| **Average B-factor** | 29.95 |  |  |  |
| macromolecules | 28.69 |  |  |  |
| solvent | 36.35 |  |  |  |

Statistics for the highest-resolution shell are shown in parentheses.

**Table S2.** Summary of the TMEM95 and IZUMO1 monoclonal antibodies

| Antigen | Antibody |  | Isotype | V-gene | D-gene | J-gene | CDR1 | CDR2 | CDR3 |
| --- | --- | --- | --- | --- | --- | --- | --- | --- | --- |
| TMEM95 | 3A01 | V<sub>H</sub> | IgG1 | IGHV5-9-4*01 | IGHD1-1*01 | IGHJ4*01 | NYVMS | EISTYGRYTFYPDSVTG | RDYYGSSSVMDY |
|  |  | V<sub>L</sub> | Kappa | IGKV4-57-1*01 |  | IGKJ1*01 | RASSSVSSSSLH | STSNLAS | QQYSGYPLT |
|  | 6B08 | V<sub>H</sub> | IgG1 | IGHV5-6*01<br>IGHV5-6-1*01 | N/A | IGHJ4*01 | TYGMS | TISFYGHTTYYPDILKG | EDYDAMDY |
|  |  | V<sub>L</sub> | Kappa | IGKV9-120*02 |  | IGKJ2*01 | RASQDIGSNLN | ATSSLDS | LQYAIFPYT |
| IZUMO1 | 4E04 | V<sub>H</sub> | IgG1 | IGHV2-6-5*01 | IGHD2-1*01<br>IGHD2-10*01<br>IGHD2-10*02 | IGHJ2*01 | DFGIS | LIWGGGNTYYNSALKS | HGRFGNTPDY |
|  |  | V<sub>L</sub> | Kappa | IGKV1-110*01 |  | IGKJ1*01 | TSGQSLVQSNGNTYLH | KVSNRFS | SQSTRFPWT |
|  | 6F02 | V<sub>H</sub> | IgG1 | IGHV3-1*02 | IGHD2-4*01<br>IGHD2-9*02 | IGHJ2*01 | SAYVWH | YIQYSGSTNYNPSLTS | AMITRGYFDY |
|  |  | V<sub>L</sub> | Kappa | IGKV14-111*01 |  | IGKJ4*01 | KASQDSNSYLS | GANRLVD | LQYDEFPFT |

**Table S3.**
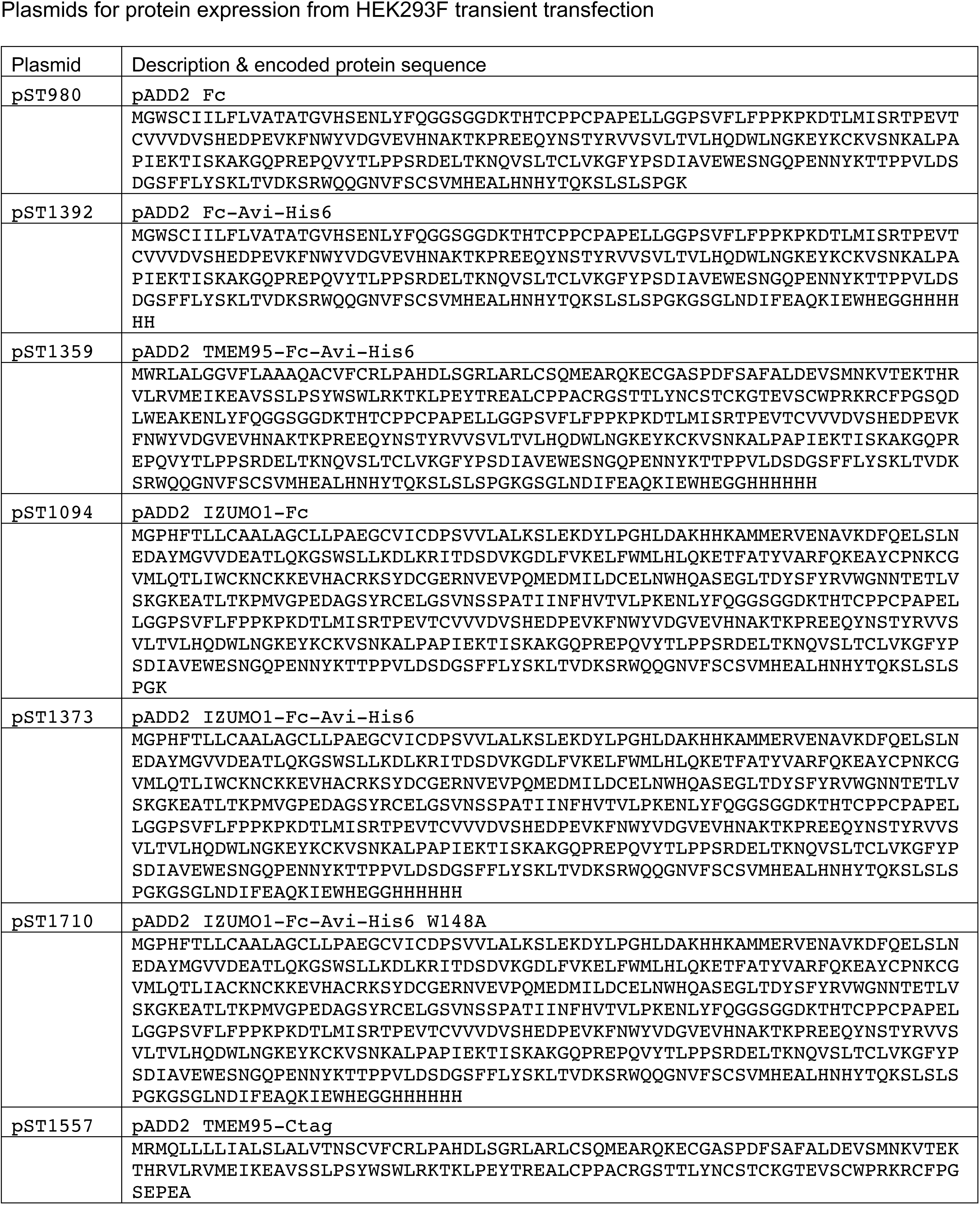
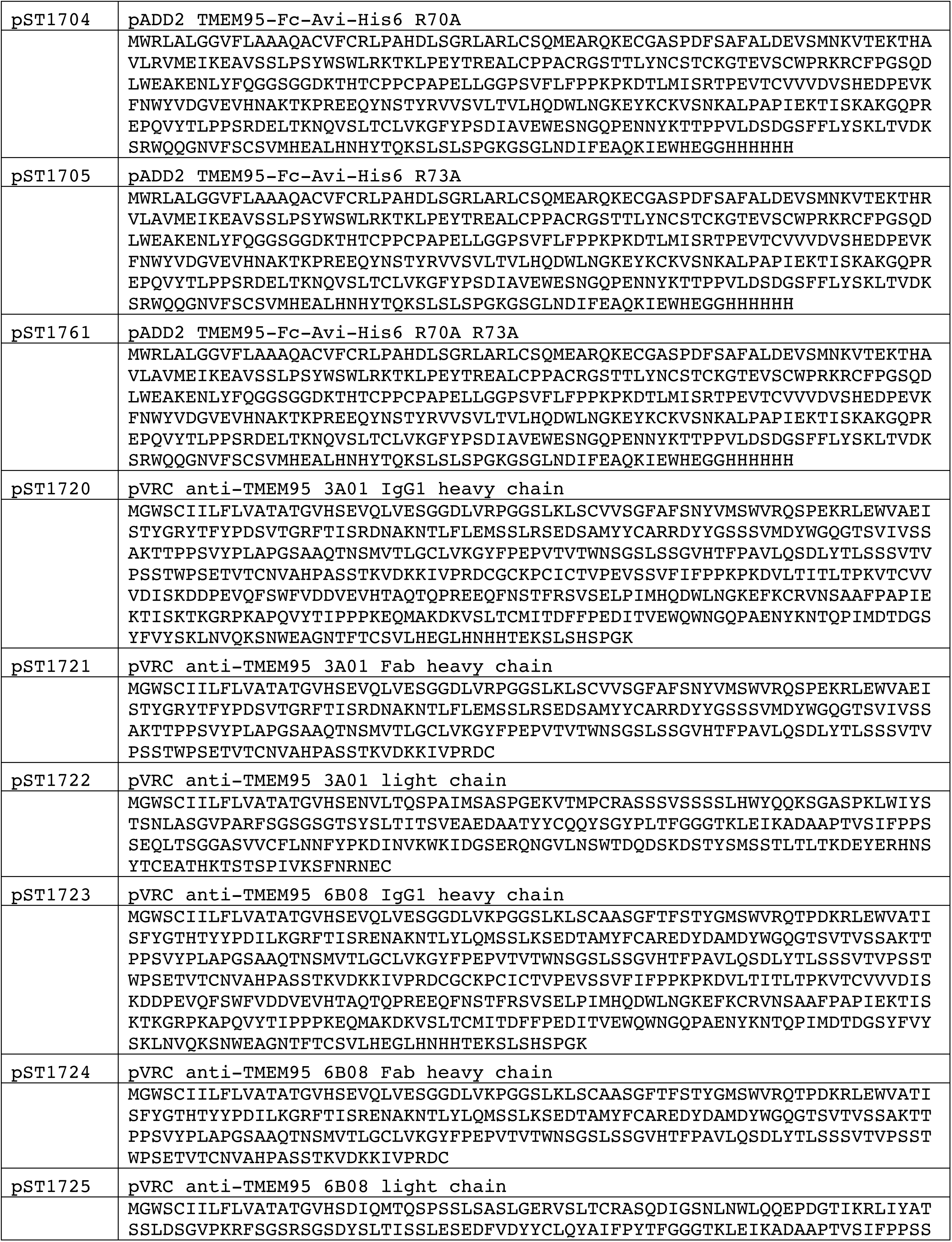
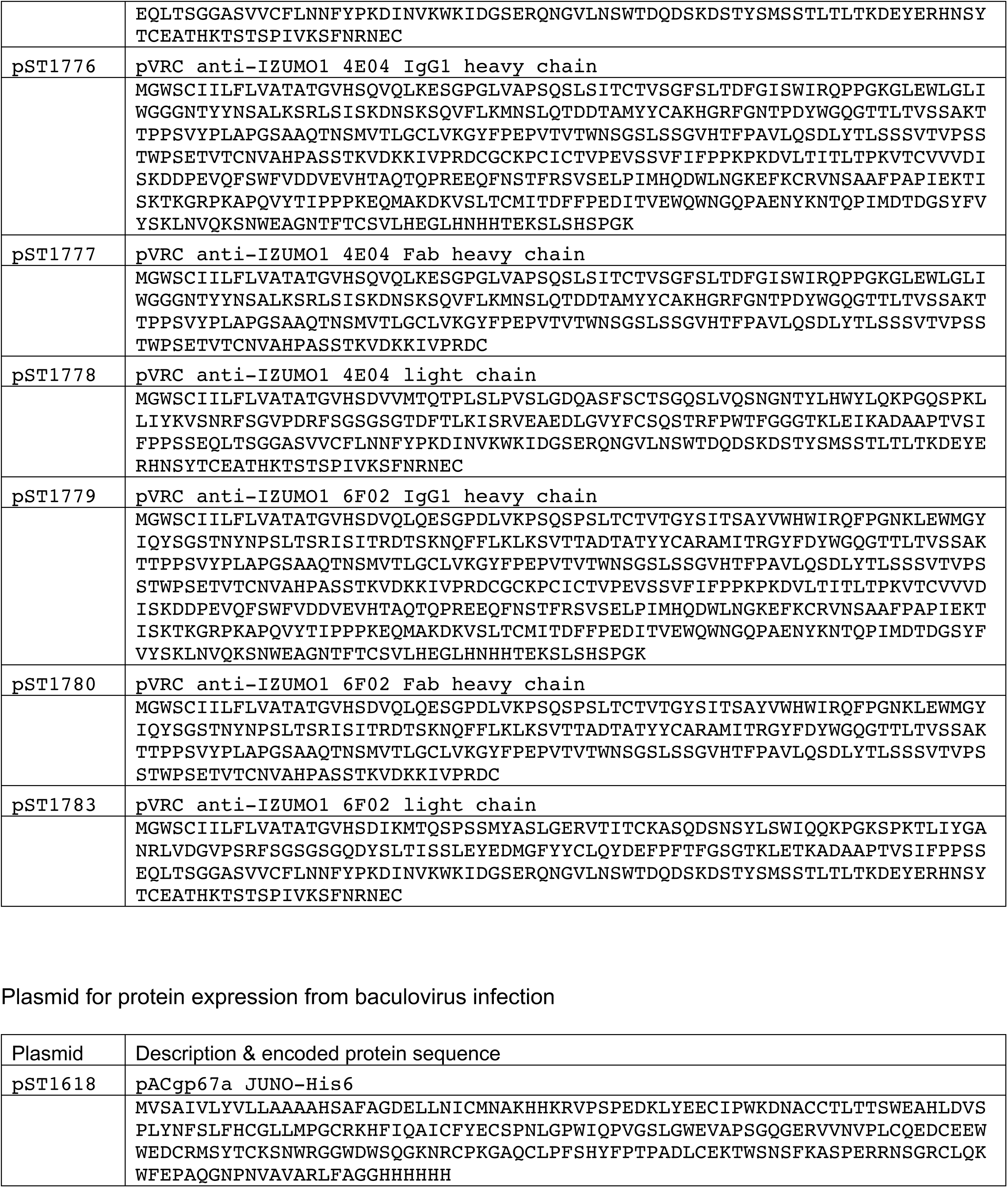
Plasmids and protein sequences used in this study

